# The stability of NPM1 oligomers regulated by acidic disordered regions controls the quality of liquid droplets

**DOI:** 10.1101/2023.01.23.525122

**Authors:** Mitsuru Okuwaki, Shin-Ichiro Ozawa, Shuhei Ebine, Motoki Juichi, Tadanobu Umeki, Kazuki Niioka, Taiyo Kikuchi, Nobutada Tanaka

## Abstract

A nucleolus is a typical membrane-less nuclear body that is formed by liquid–liquid phase separation (LLPS) of its components. A major component that drives LLPS in the nucleolus is nucleophosmin (NPM1). The oligomer formation and inter-oligomer interactions of NPM1 are suggested to cooperatively contribute to the induction of LLPS. However, the molecular mechanism of how the quality of the liquid droplets formed by NPM1 is regulated is currently not well understood. In this manuscript, we revealed the regulatory mechanism of NPM1 oligomer formation and its relationship with the ability to form liquid droplets. Molecular dynamics simulations and mutant protein analyses suggest that the acidic amino acids in the N-terminal and central disordered regions of NPM1 disturb the key interactions between monomers. We also demonstrate that mutants with attenuated oligomer stability form liquid droplets as do the wild-type; the fluidity of the formed liquid droplets was greater than that of the wild-type. These results suggest that the stability of NPM1 oligomers is a critical determinant of liquid droplet quality. Furthermore, we observed that when the net negative charges in the acidic disordered regions were increased by phosphomimetic mutations at Ser125, the NPM1 oligomer stability decreased, which increased the fluidity of the liquid droplets. Our results provide a novel mechanistic insight into how nucleolar dynamics are regulated during the cell cycle.

## Introduction

NPM1/nucleophosmin is a phosphoprotein that is mainly localized in the nucleolus. NPM1 positively regulates the transcription of precursor ribosome RNA (pre-rRNA) as a histone chaperone [1, 2] and the processing of pre-rRNA [3, 4]. In addition, NPM1 regulates the functions of the p53 pathway by direct binding with p53 or a tumor suppressor Arf [3, 5, 6]. NPM1 also localizes to the centrosome and regulates the centrosome duplication, and NPM1 depletion induces uncontrolled centrosome duplication, resulting in chromosome instability [7, 8]. Given these multiple functionalities, *NPM1* is an essential gene [8].

NPM1 consists of three structural regions: the oligomerization domain (amino acids 14–117), the C-terminal globular domain (amino acids 243–294), and the intrinsically disordered region (IDR) between the N- and C-terminal structured domains [9]. The N-terminal 15 amino acid region is also likely to be disordered. The oligomerization domain forms a stable pentamer, and two pentamers are further assembled into a decamer in solution [10, 11]. The IDR is divided into two regions, an acidic and a basic amino acid-rich regions [12], which are hereafter referred to as aIDR and bIDR, respectively. The C-terminal globular domain consists of three alpha helices and plays a key role in recognizing RNA with the help of the bIDR [13]. RNA recognition by this domain is required for the nucleolar localization of NPM1 [12]. Ser4 in the N-terminal disordered region, Ser125 in the aIDR, and Thr199, Thr219, Thr234, and Thr237 in the bIDR, are phosphorylated; phosphorylation of the latter four threonine residues during mitosis inactivates the RNA binding activity of NPM1 [14–16]. Dissociation of NPM1 from RNA has been suggested to cause nucleolar disassembly during mitosis [16, 17]. Ser125 is likely to be constitutively phosphorylated after translation, and this phosphorylation regulates the chaperone activity of NPM1 [15], although its biological significance remains elusive.

NPM1 is generally translated from the first AUG codon on its mRNA, as are most mRNAs. It has been reported that the 2nd, 3rd, and 4th AUG codons are also used for the initiation of translation; as a result, NPM1 proteins that are translated from the 1st, 5th, 7th, and 9th methionine (Met) are expressed in cells [18–20]. This initiation of translation from the non-first AUG codons is likely to be an example of alternative translation initiation due to the leaky scanning of the ribosome [21]. Interestingly, it was reported that NPM1 translated from the 7th Met (M7-NPM1) forms a more stable oligomer than normal NPM1 translated from the 1st Met [19, 22], although the biological significance of the expression of M7-NPM1 and how the initial six amino acids lacking in M7-NPM1 affect the oligomer formation of NPM1 are unknown.

The nucleolus, a membrane-less nuclear body, is the site of ribosome biogenesis [23]. Ribosome biogenesis is an energy-consuming process that includes transcription of pre-rRNA with approximately 12,000 bases and its stepwise processing to produce three mature rRNAs (18S, 5.8S, and 28S rRNAs). During transcription by RNA polymerase I and during the processing of pre-rRNA, ribosomal proteins are deposited on pre-rRNAs to assemble mature ribosomes. A variety of pre-rRNA processing and ribosome assembly factors that are localized in the nucleoli are required for the stepwise processing and assembly of ribosomes [24].

The nucleolus is not surrounded by a membrane, and it has long been a mystery how the nucleolus is formed in the nucleus. Recent research has demonstrated that the biophysical properties of protein, RNA, and DNA play key roles in the formation of membrane-less cellular bodies, including nucleoli [25]. NPM1 has been reported to produce homotypic liquid droplets by liquid–liquid phase separation (LLPS) under certain conditions [26–28]. Monomer–monomer interactions that form the NPM1 pentamers, as well as inter-pentamer interactions through the interaction between aIDR and bIDR, are likely to be required for LLPS. In addition, NPM1 forms heterotypic liquid droplets by associating with arginine-rich proteins or rRNAs [26, 29]. We demonstrated that nucleolar morphology, particularly the outer layer of the nucleolus referred to as granular components (GC), was clearly changed upon NPM1 depletion owing to partial diffusion of the large ribosome subunit precursors throughout the nucleus [4]. These results strongly suggest that LLPS is a driving force for the formation of nucleoli and that NPM1 is a scaffold component that forms the GC region of the nucleolus.

In this manuscript, we revealed the regulatory mechanisms for NPM1 pentamer formation and the significance of the regulation of the NPM1 oligomers in the assembly and disassembly of the nucleolus during the cell cycle. Our results suggest that fluctuations in NPM1 oligomer stability are closely related to nucleolar assembly and disassembly during the cell cycle.

## Results

### NPM1 oligomer stability is regulated by acidic amino acids in the IDRs

The mRNA that codes for NPM1 has several AUG codons at the 5’ region (Figures 1A and B) and the first AUG is not always used for the initiation of translation, which can result in the expression of N-terminally truncated products. The NPM1 protein (M7-NPM1) that is translated from the third AUG codon (7th Met) was demonstrated to be expressed in the liver [19]. Furthermore, a previous report demonstrated that M7-NPM1 forms a more stable oligomer than NPM1 that was translated from the first AUG [22]. We first sought to clarify how alternative initiation of translation of NPM1 affects the stability of the NPM1 oligomer. Consistent with the previous results, NPM1 with a molecular weight of 37 kDa was partially detected as high molecular weight oligomers, even in SDS-PAGE sample buffer that contained 1% SDS when the protein was loaded onto the gel without heating before electrophoresis (Figure 1B, lane 1). Based on previous native PAGE and gel filtration analyses, the oligomer bands was demonstrated to correspond to the decamer [30]. When the sample was incubated at 95°C before electrophoresis, the high molecular weight oligomer was mostly disrupted (compare Figure 1B, lanes 1 and 9). The NPM1 proteins that was translated with the second or third AUGs (M5-NPM1 or M7-NPM1) were resistant to SDS-PAGE sample buffer containing 1% SDS, and high molecular weight oligomers were detected (lanes 2 and 3). Upon heat denaturation, the oligomer was mostly disrupted, but small numbers of oligomers were still detected (lanes 10 and 11). In addition, the amount of M7-NPM1 oligomers was greater than that of M5-NPM1 oligomers. These results suggest that the first four and additional two amino acids contribute to attenuating the stability of the NPM1 oligomer. The N-terminal peptide contains acidic amino acids at the 2nd, 3rd, and 6th positions (Figure 1B). This sequence prompted us to examine whether the acidic amino acids affect the stability of the NPM1 oligomer. To examine this possibility, we prepared point mutant proteins where 2nd to 7th amino acids were individually or concomitantly substituted with alanine (NPM1-6A, 23A, 236A, 2346A, and 234567A, see Figure 1B top panel). We found that substitutions of the 2nd, 3rd, and 6th acidic amino acids with alanine augmented the stability of the NPM1 oligomer, and the high molecular weight bands of 23A and 6A were much clearer than that of the wild-type NPM1 regardless of whether the samples were heated or not before loading onto SDS-PAGE gels. In addition, when the 2nd Glu (E), 3rd Asp (D), and 6th Asp (D) were concomitantly substituted with alanine (236A), the stability of the oligomer of mutant (236A) was similar to that of M7-NPM1, whereas 2346A and 234567A exhibited similar oligomer stability to the 236A mutant (lanes 6–8 and 14–16). These results indicate that the acidic property of the N-terminal disordered region attenuates the stability of NPM1 oligomers.

**Figure 1.**
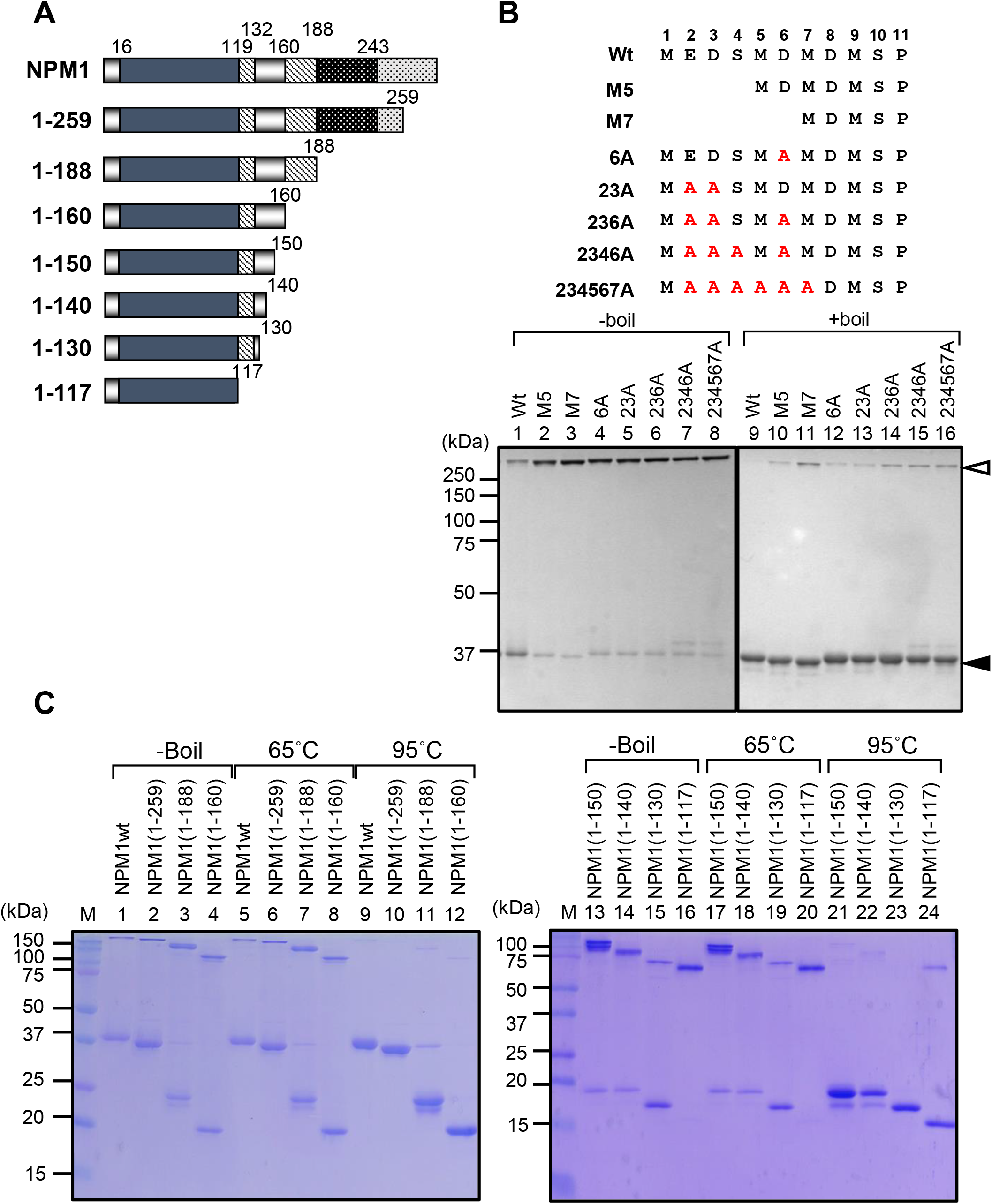
The N-terminal and central acidic regions of NPM1 destabilize SDS-resistant oligomers. A. Schematic representations of NPM1 mutants. NPM1 wild-type and mutants used in Figure 1 are schematically represented. The gray region (amino acids 16–119) is the oligomerization domain, shaded boxes (amino acids 119–132 and 160–188) are the acidic regions, dark filled region (amino acids 243–294) is the folded RNA binding domain. B. Effects of N-terminal deletion and point mutations on the oligomer stability of NPM1. The wild-type and mutant NPM1 proteins (500 ng) indicated at the top of each lane were incubated in the buffer containing 1% SDS and loaded on 10% SDS-PAGE without (lanes 1–8) or with heating at 95°C for 5 min (lanes 9–16). The proteins were visualized with CBB-staining. The positions of NPM1 monomers and oligomers are indicated by filled and blank arrowheads, respectively, at the right. The positions of molecular weight markers are shown at the left of the panel. B. Effects of central acidic regions on the oligomer stability. The wild-type and mutant NPM1 proteins (500 ng) indicated at the top of each lane were incubated in the buffer containing 1% SDS and loaded on 12.5% SDS-PAGE without heating (lanes 1–4 and 13–16) or heating at 65°C (lanes 5–8 and 17–20) or 95°C (lanes 9–12 and 21–24) for 5 min.

Given that NPM1 contains two highly acidic stretches in its central disordered region (Figure 1A), we questioned whether these acidic stretches also play roles in regulating the stability of NPM1 oligomers as does the N-terminal region. To test this possibility, we prepared a series of C-terminally truncated mutants; NPM1(1–259), NPM1(1–188), NPM1(1–160), NPM1(1–150), NPM1(1–140), NPM1(1–130), and NPM1(1–117) (Figure 1A). The proteins were separated by SDS-PAGE before or after heating at 65°C or 95°C and visualized by CBB staining (Figure 1C). The results demonstrated that NPM1(1–117), which lacks the central acidic stretches, was mostly resistant to SDS, and oligomer bands at 75 kDa were detected when the samples were loaded onto SDS-PAGE gels without heating or by heating at 65°C (lanes 16 and 20). Moreover, the oligomer band was detected even after heating at 95°C (lane 24). In addition, the oligomer bands for NPM1(1–259), NPM1(1–188), NPM1(1–160), NPM1(1– 150), NPM1(1–140), and NPM(1–130) were similarly detected when the samples were loaded onto SDS-PAGE gels before or after heating at 65°C as was full-length NPM1 (lanes 1–8, 13–15, and 17–19); however, their oligomer bands were eliminated after heating at 95°C (lanes 9–12 and 21–23). These results indicate that the acidic regions in the central IDRs also attenuate the stability of the oligomers, and the region between amino acids 120 and 130 of NPM1 is sufficient to destabilize the NPM1 oligomers.

### Molecular dynamics simulation of the interactions between acidic disordered regions and the oligomerization domain

Because the N-terminal region (amino acids 1 to 13) and the central acidic regions of NPM1 are intrinsically disordered and these regions in the crystal structure cannot be detected [10, 11], it is difficult to clarify the molecular mechanism by which these disordered regions regulate the stability of the NPM1 oligomer. To gain insight into this mechanism, we performed molecular dynamics (MD) simulations starting from the NPM1 pentamer structures with (NPM1[1–118] and NPM1[14–130]) or without (NPM1[14–118]) the acidic IDRs. Upon close examination of the crystal structure of NPM1, the amino acids involved in monomer–monomer interactions were identified (Figure 2A). From the MD trajectories, we obtained 12,500 solution structures for each NPM1 pentamer and then enumerated the number of contacts (hydrogen-bonding, ionic, π–π stacking, and π–cation interactions) in these structures. Figure 2B shows the structures of the NPM1 pentamer containing the N-terminal (M1–A118) or central acidic (P14–E130) regions. The disordered acidic regions (represented by green tubes) move freely with some preference for the oligomerization domain. The interactions between the oligomerization domain and the IDRs were estimated and are shown using a heatmap (Figure 2C). The residues in the oligomerization domain (the left column in Figure 2C) can contact any residue in the acidic IDRs (the top row of Figure 2C). Some residues, such as Gln15, Glu37, Asn38, and Glu39, are located in the solvent-accessible region and appear to be in contact with the acidic IDRs; others (Tyr17, Lys32, Asp34, Asp36, His40, Tyr67, and Glu68) are involved in monomer–monomer interactions. Previous crystal structures suggested that Tyr67 plays a key role in connecting two monomers as a latch (Figure 2A) [11]. In the NPM1 pentamer, Tyr67, located at the tip of the loop between β4 and β5, is surrounded by three aromatic amino acids, Tyr17, His40, and His115, from an adjacent subunit, presumably to make contact by π–π stacking [11] (Figure 2A). Tyr67 may also make contact with Asp34 or Asp36 from an adjacent subunit via hydrogen bonding. The key amino acids involved in monomer–monomer interactions are located at a site that is accessible to the N-terminal acidic amino acids. We also calculated contact frequencies between amino acids that are involved in monomer–monomer interactions, i.e., Lys32 and Glu68, Asp34 and Tyr67, Asp36 and Tyr67, Tyr17 and Tyr67, His40 and Tyr67, and Tyr67 and His115, in the presence (M1–A118 and P14–E130) or absence (P14–A118) of acidic disordered regions (Figure 2D). Although the interaction frequency between Tyr17 and Tyr67 was not clearly affected by the presence of N-terminal and central acidic regions, the others were increased by removing the acidic disordered regions. In particular, the interaction frequency between H40 and Y67 was relatively high (0.439) in the absence of acidic regions (P14–A118), whereas it decreased upon adding the N-terminal (0.341 for M1–A118) or central (0.338 for P14–E130) acidic disordered regions.

**Figure 2.**
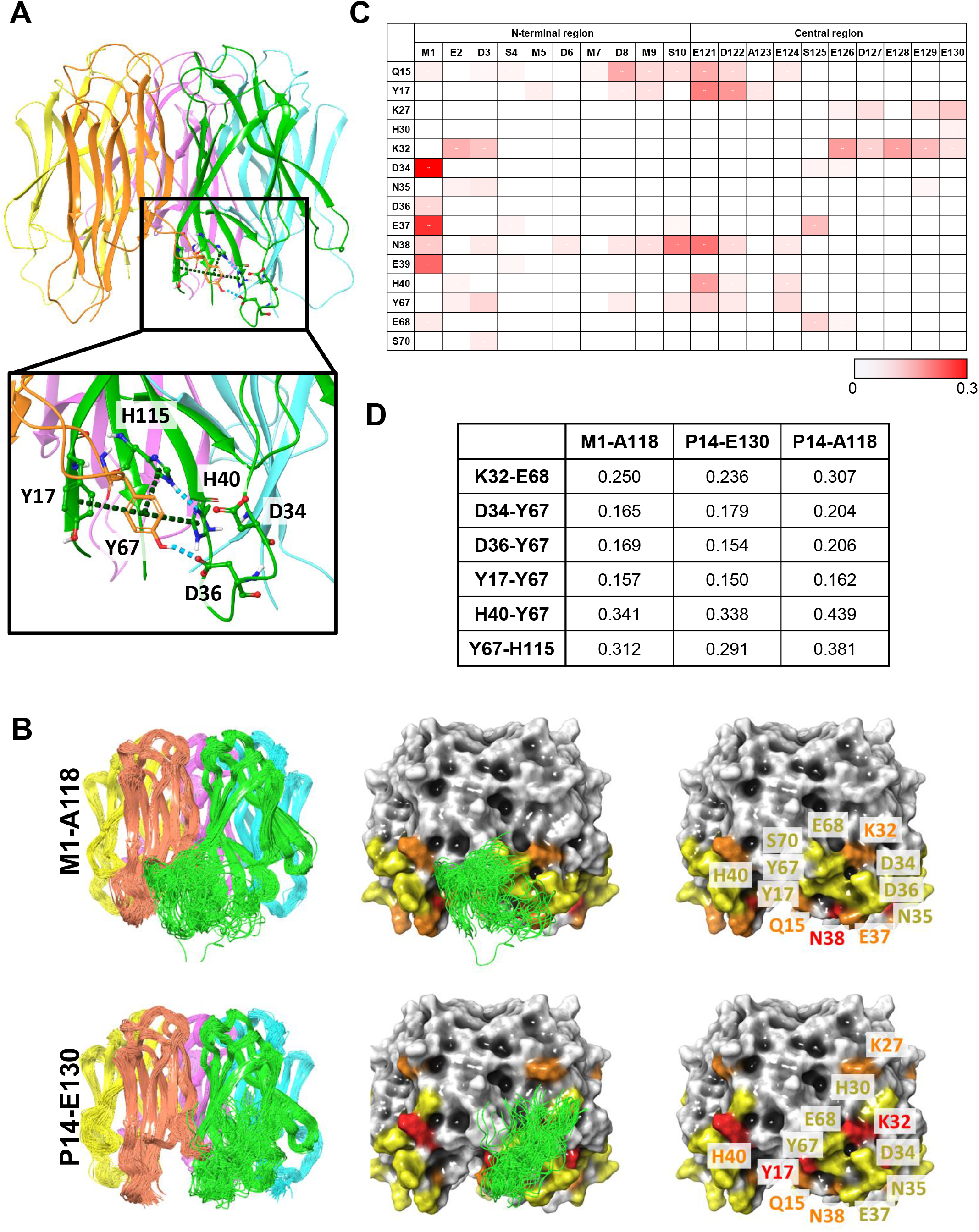
MD simulations for the interactions between the aIDRs and the oligomerization domain of NPM1. (A) Crystal structure of the NPM1 pentamer (PDB code: 5EHD). The pentamer structure and the interaction network around Tyr67 (top and bottom panels) are shown. Hydrogen bonds and aromatic interactions are shown as cyan and green dotted lines, respectively. (B) The positions of the aIDRs in the representative structures of the MD simulations. Successive 100 structures are selected from 2,500 snapshots along 50 ns simulation time. The acidic IDRs are shown as green tubes,, and yellow to red shading on the surface of the oligomerization domain indicates residues with increasing frequency of contact with the acidic IDRs (yellow: 0.01 to 0.15, orange: 0.15 to 0.30, and red: higher than 0.30). (C) Frequency heatmap of the contacts between the oligomerization domain and the N-terminal and central IDRs. The residues in the oligomerization domain (left column) in contact with any residues in the N-terminal and central IDRs (top horizontal row) (frequency higher than 0.01) are shown. (D) Effects of the acidic IDRs on the frequency of the monomer–monomer interactions within the oligomerization domain. The NPM1 pentamer with N- (M1–A118) or central (P14–E130) acidic regions, or without IDRs (P14–A118) were subjected to MD simulations as in C and the interaction frequency between key residues involved in the monomer–monomer interactions (shown in the left column) were calculated.

To validate the results that were obtained from the MD simulations, we prepared point mutant proteins and determined the stability of their oligomers (Figure 3A). To evaluate the importance of potential interactions between Lys32 and Glu68, Lys32 was substituted with alanine (K32A). NPM1, M7-NPM1, NPM1-K32A, and M7-NPM1-K32A were heated at 95°C for 0, 1, and 5 min, followed by SDS-PAGE and CBB staining (Figure 3B). Consistent with the results shown in Figure 1, the high molecular weight bands of M7-NPM1 were greater compared to those of wild-type NPM1 and the M7-NPM1 oligomer bands were still detected after heating the sample at 95°C for 5 min (lane 6). Similar results were obtained when NPM1-K32A and M7-NPM1-K32A were examined (lanes 7–12). To analyze the effect of the mutation on oligomer formation, we prepared proteins that contained only the oligomerization domain (amino acids 7–117, Figure 3A). The samples were analyzed by SDS-PAGE after heating at 95°C for 1, 2, 3, 4, or 5 min (Figure 3C). The results demonstrated that the K32A mutation affects neither oligomer formation nor oligomer stability, and the oligomer bands were detected as they were with NPM1(7–117).

**Figure 3.**
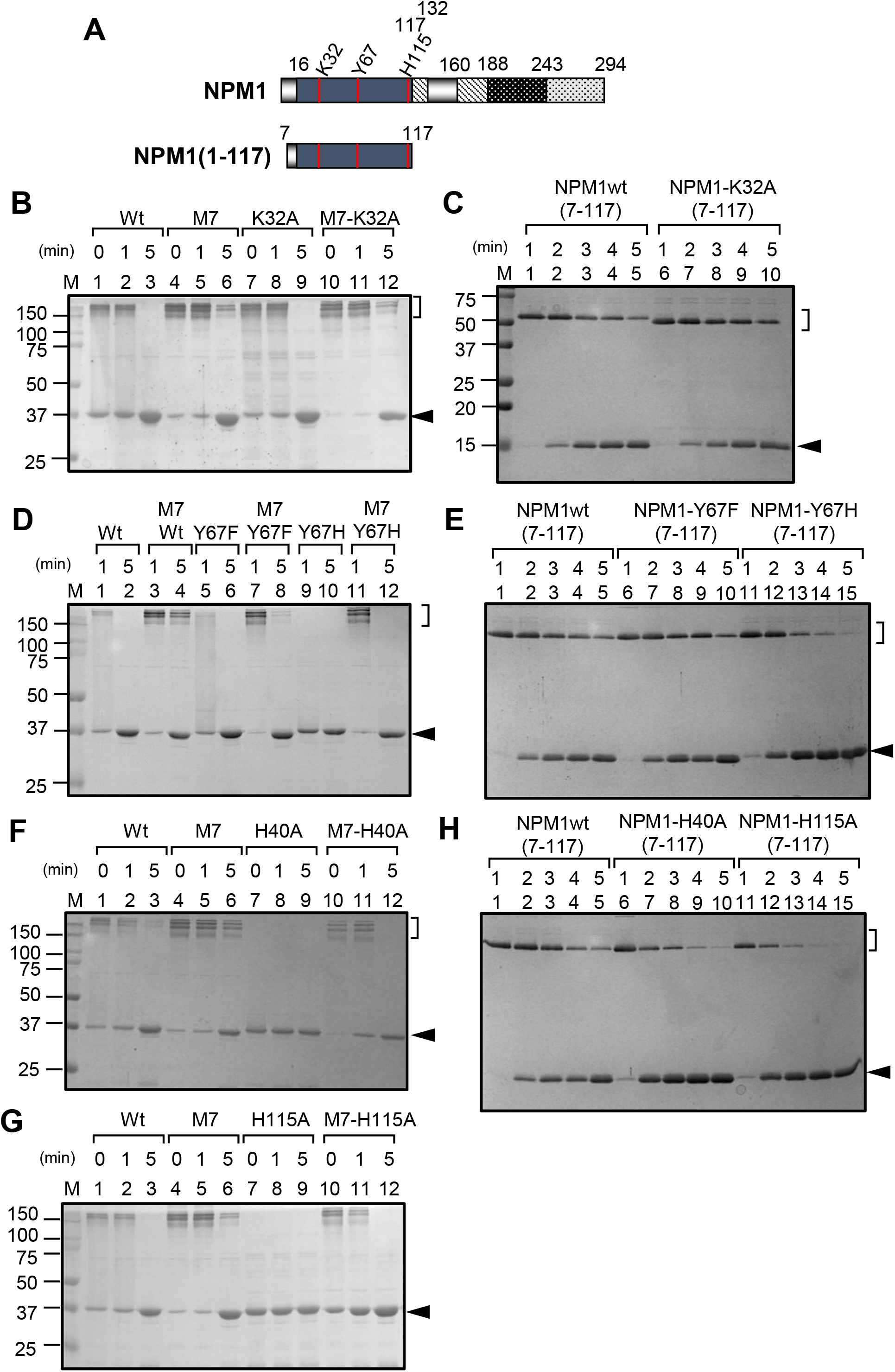
Effects of point mutations in the oligomerization domain on the stability of the NPM1 oligomers. A. Schematic representation of point mutant NPM1 and NPM1 oligomerization domain. Full-length NPM1 and NPM1(7–117) are shown and the positions of amino acids mutated are shown with red bars. B, D, F and G. Effects of mutations on the NPM1 oligomer stability. Wild-type and NPM1-K32A (B), NPM1-Y67H or Y67F (D), NPM1-H40A (F), and NPM1-H115A (G), and M7-NPM1 and its mutants were mixed with SDS-PAGE loading dye and separated by SDS-PAGE before or after 1 min or 5 min heating at 95°C as shown at the top of each panel. C, E, H. Effects of mutations on the oligomer stability of the NPM1 oligomerization domain. NPM1(7–117) and its mutants were separated by 12.5% SDS-PAGE after 1, 2, 3, 4, or 5 min heating at 95°C as shown at the top of each lane. Proteins were visualized by CBB staining and positions of monomers and oligomers are indicated at the right of the panels.

Next, to examine the interactions between Tyr67 and Asp34 or Asp36, we prepared NPM1 proteins in which Tyr67 was substituted with phenylalanine (Y67F) or histidine (Y67H). We chose phenylalanine to substitute for Tyr67 to specifically disturb the hydrogen bonding while maintaining the aromatic nature, whereas we chose histidine because, in NPM3, an NPM family member, the position corresponding to the Tyr67 in NPM1 is histidine [30]. We found that the oligomer band of the Y67F mutant after 1 min heating was slightly less in the intensity than that of wild-type NPM1, whereas both wild-type and Y67F oligomer bands were disrupted by 5 min of heating (Figure 3D lanes 1, 2, 5, and 6). Similarly, the oligomer bands for M7-NPM1-Y67F were slightly lower in intensity than those for M7-NPM1 (lanes 3, 4, 7, and 8). Furthermore, high-molecular weight oligomer bands for NPM1-Y67H were not detected even after 1 min of heating (lanes 9 and 10). High-molecular weight bands for M7-NPM1-Y67H were clearly detected after feating for 1 min and essentially disappeared after heating for 5 min (lanes 11 and 12). In addition, we compared the stability of oligomers for Tyr67 mutants using proteins harboring amino acids 7–117. The stability of oligomers for NPM1(7–117)-Y67F was similar to that of NPM1(7–117) (Figure 3E, lanes 1–10). In contrast, NPM1(7–117)-Y67H exhibited less oligomer stability than NPM1(7–117) (lanes 11–15). These results suggest that stacking interactions between Tyr67 and His40 or His115 play a major role, while the ionic interactions between Tyr67 and Glu34 or Glu36 play a minor role in the formation of NPM1 oligomers.

To confirm the significance of His40 and His115 in the formation of NPM1 oligomers, we prepared mutants wherein His40 or His115 was replaced with alanine (H40A or H115A). We observed that NPM1-H40A and NPM1-H115A produced no high molecular weight oligomer bands detectable by SDS-PAGE analysis even when the proteins were analyzed without heating (lanes 7 of Figures 3F and 3G). When M7-NPM1-H40A and M7-NPM1-H115A were examined, both showed high-molecular-weight oligomer bands when samples were loaded on SDS-PAGE without heating or after heating for 1 minute (lanes 10 and 11 in Figures 3F and G), although the bands were almost eliminated after 5 min of heating (lanes 12). Furthermore, NPM1(7–117)-H40A and NPM1(7–117)-H115A exhibited a much lower ability to form oligomers than NPM1(7–117) (Figure 3H). From these results, we conclude that the interactions between Tyr67 and His40 as well as Tyr67 and His115 play crucial roles in the formation of NPM1 oligomers. Considering that the M7-NPM1 mutant proteins (Y67H, H40A, and H115A) form oligomers, the mutations Y67H, H40A, and H115A did not completely disrupt the oligomer formation.

To gain further insight into the function of the N-terminal and central acidic amino acids in the regulation of NPM1 oligomer stability, the MD simulation data shown in Figure 2 were analyzed. Figure 4 shows representative snapshots, including contacts between the oligomerization domain (Tyr67 or His40) and the acidic amino acids in the IDRs (Glu2 or Asp124). Hydrogen-bonding and ionic interactions between the acidic amino acids in the IDRs and Tyr67 or His40 may disrupt the aromatic interaction network formed by His40, Tyr67, and His115, resulting in destabilization of the NPM1 oligomer structure.

**Figure 4.**
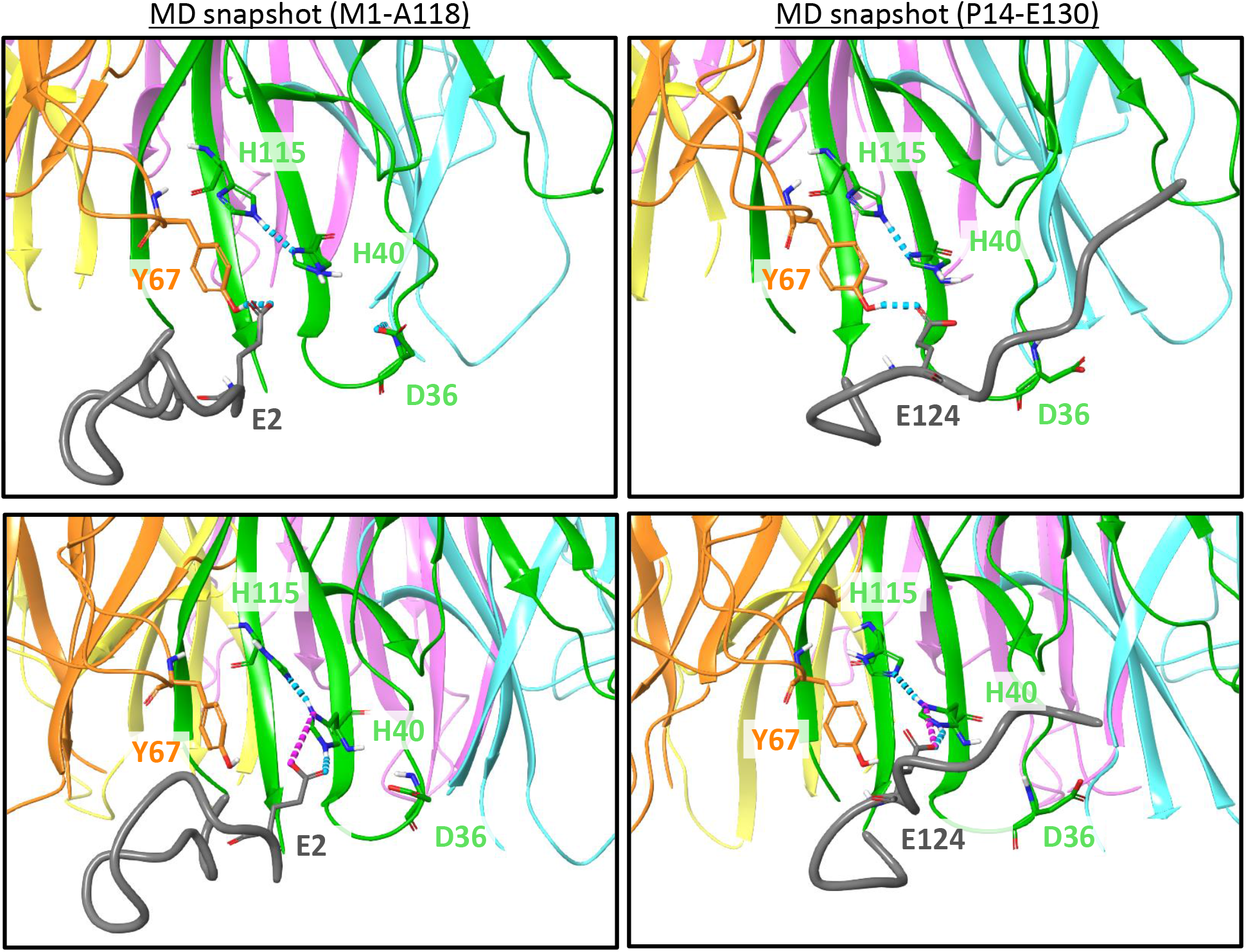
Snapshots of the interaction between the N-terminal or central acidic regions and the oligomerization domain. The snapshots for the NPM1 pentamer including the N-terminal and central IDRs are shown (left and right panels, respectively). The top panels are snapshots showing the interactions between Tyr67 and the aIDRs, whereas the bottom panels are snapshots showing the interactions between His40 and the acidic IDRs. The acidic IDRs are shown as gray tubes. Hydrogen bonds and salt bridges are shown as cyan and magenta dotted lines, respectively.

### NPM1 oligomer stability regulates liquid droplet fluidity

NPM1 is a major nucleolar protein, and it plays a critical role in the formation of the nucleolus via its ability to form liquid droplets. The formation of liquid droplet by NPM1 requires oligomer formation and inter-oligomer interactions [27]. As shown in Figures 1–4, the acidic amino acids in the N-terminal region and the aIDR of NPM1 may contact the oligomerization domain and decrease the stability of the oligomers. We questioned whether the stability of NPM1 oligomers is associated with the process of liquid droplet formation. NPM1 and M7-NPM1 were labeled with IC3-OSu dye, and images of the liquid droplets were obtained by confocal microscopy with bright-field and fluorescence using a 560 nm-laser (Figures 5A and B). Homotypic LLPS induced the formation of liquid droplets by NPM1 as previously reported [27] in ithe buffer containing 150 mM salt (Figure 5A, left panels). In addition, M7-NPM1 formed liquid droplets, as did wild-type NPM1. We observed that the liquid droplets with M7-NPM1 tended to be larger than those with NPM1. When RNAs purified from 293T cells that contained rRNAs were added to the sample, the NPM1 wild-type formed larger droplets that possibly include RNAs, indicating that NPM1 and RNA cooperatively formed heterotypic liquid droplets (Figure 5A, right panels). Heterotypic droplets were also formed with M7-NPM1 and RNA. Next, we examined liquid droplet formation with NPM1-H40A, M7-NPM1-H40A, NPM1-H115A, and M7-NPM1-H115A. As shown in Figure 3, NPM1-H40A and NPM1-H115A exhibited lower oligomer stability in the presence of SDS-PAGE loading buffer containing 1% SDS. Despite the lower oligomer stability of NPM1-H40A and NPM1-H115A, these proteins formed liquid droplets as did wild-type NPM1 (Figure 5B). Although M7-NPM1-H40A and M7-NPM1-H115A also exhibited higher oligomer stability than NPM1-H40A and NPM1-H115A, we did not observe clear differences in droplet numbers and sizes (Figure 5B).

**Figure 5.**
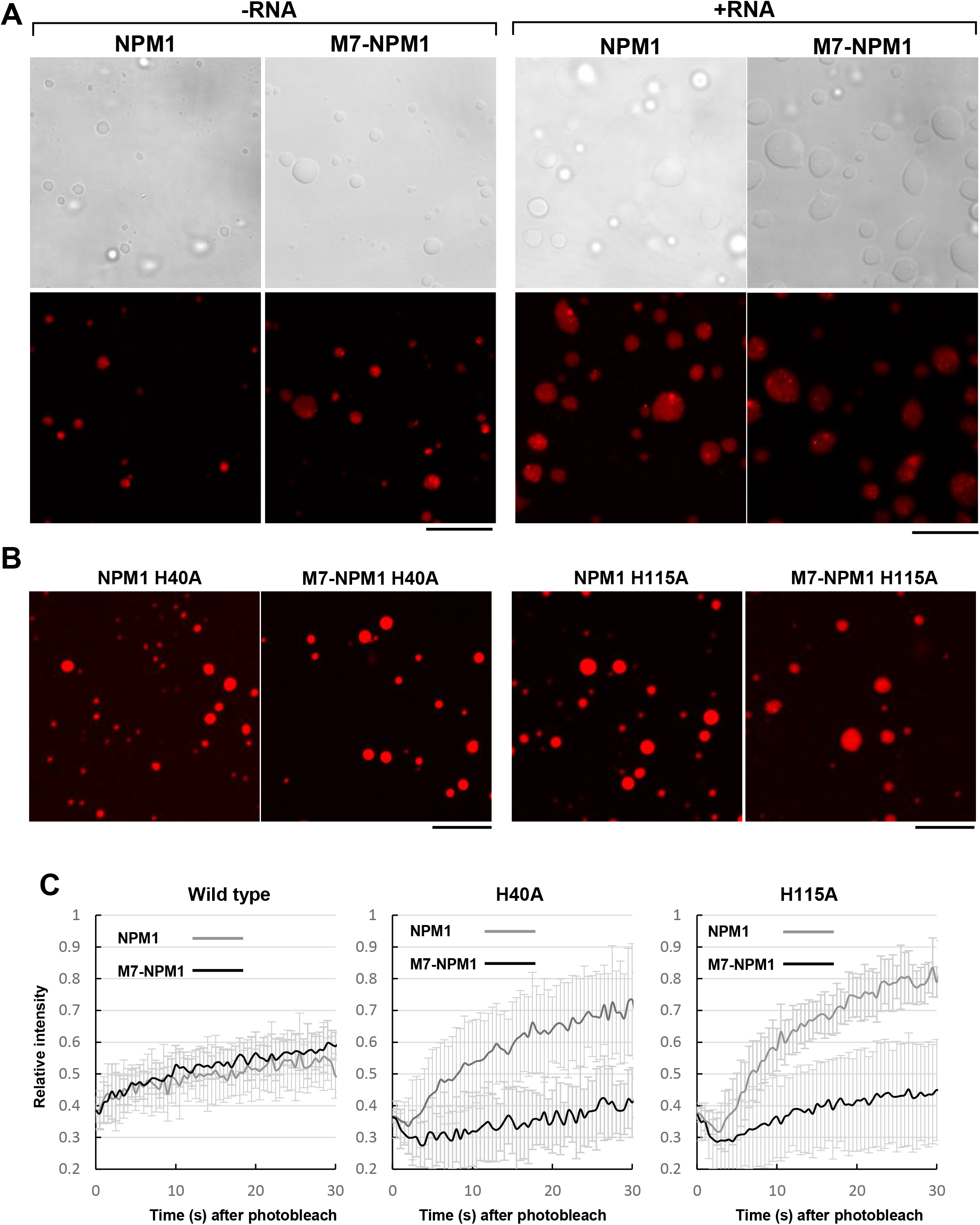
The close association between oligomer stability and the ability of NPM1 to form liquid droplets. A. Liquid droplet formation by NPM1 and M7-NPM1. The liquid droplets were formed with IC3-OSu-labeled NPM1 or M7-NPM1 (10 μM) in the absence or presence of 1 μg of total RNA extracted from 293T cells in the buffer containing 8% Ficoll PM70 and 150 mM NaCl. Bright-field and fluorescent images (top and bottom panels, respectively) are shown for each sample. Bars at the right bottom of each panel indicate 10 μm. B. Effects of point mutations at H40 and H115 on the liquid droplet formation ability of NPM1. Liquid droplets were formed with NPM1-H40A, M7-NPM1-H115A, and M7-NPM1-H115A labeled with IC3-OSu (15 μM) as in A. C. FRAP analyses of NPM1 liquid droplets. Liquid droplets were formed with 20 μM NPM1 in the buffer containing 2% Ficoll PM70 and 150 mM NaCl. FRAP analyses were performed as described in Materials and methods. The samples were prepared in a 384-well glass-based plate and the experiments were started 10 min after sample preparation. For all samples, 10 droplets were examined and error bars indicate ±SD. For each mutation, the recovery curves for NPM1 and M7-NPM1 are shown with gray or black lines, respectively.

To quantitatively analyze the quality of the droplets formed by NPM1 and M7-NPM1 mutants, fluorescence recovery after photobleaching (FRAP) assays were performed (Figure 5C). The fluorescence in the central region of the droplets was photobleached with a 561 nm-laser until the intensities dropped to less than 40% of the initial intensity before photobleaching, and the fluorescence intensities of the bleached areas were measured every 0.5 sec for 30 sec. The recovery rates for the NPM1 wild-type were relatively slow, and the intensities recovered to only 60% of the initial intensity over a period of 30 sec. We also observed that M7-NPM1 exhibited similar recovery rates, indicating that NPM1 and M7-NPM1 formed liquid droplets with similar fluidity under the assay conditions employed here. In addition, we observed that the recovery rates for NPM1-H40A and NPM1-H115A were higher than that for wild-type NPM1, and the intensities recovered to 70%–80% of the initial intensity within 30 sec. Interestingly, we observed that M7-NPM1-H40A and M7-NPM1-H115A exhibited lower recovery rates than the NPM1-H40A and NPM1-H115A mutants. These results indicate that the stability of NPM1 oligomers determines liquid droplet fluidity.

### Effects of interphase and mitotic phosphorylation on NPM1 oligomer stability

NPM1 is known to be phosphorylated at Ser125 during interphase [15, 17] and at Ser4 [14], Thr199, Thr219, Thr234, and Thr237 during mitosis [16]. Ser4 and Ser125 are located at the N-terminal and aIDR regions, respectively, and both are involved in the destabilization of NPM1 oligomers (Figure 1). In addition, Thr199, Thr219, Thr234, and Thr237 are located in the bIDR. Although the biological significance of Ser4 and Ser125 phosphorylation is currently unclear, the phosphorylation of Thr199, Thr219, Thr234, and Thr237 decreases the RNA binding activity of NPM1 [12, 16], which is believed to contribute to nucleolar disassembly during mitosis. The increase in the net negative charges on the IDRs by phosphorylation may enhance the oligomer destabilization activity of the disordered regions. To test this, the phosphomimetic mutants S4D, S125D, and T4sD (Thr199, Thr219, Thr234, and Thr237 are replaced with aspartic acids) were prepared, and the stability of their oligomers was examined (Figure 6). Purified proteins were mixed with SDS-loading buffer and incubated at room temperature, 65°C, 75°C, 85°C, and 95°C for 5 min followed by SDS-PAGE analysis. Approximately half of the wild-type NPM1 was detected as a monomer when the sample was incubated at room temperature before electrophoresis, and the amount of monomer gradually increased as the incubation temperature increased (Figures 6A–C, lanes 1–5). A similar increase in the amount of monomers was observed for all phosphomimetic mutants, although the amount of monomers abserved after incubation at room temperature for NPM1-T4sD and NPM1-S125D were clearly more than that of wild-type (Figures 6B, C, lanes 6). NPM1-S4D monomer was also slightly more than the wild-type NPM1 monomer, although the difference between S4D and wild-type was not extremely clear (Figure 6A). These results indicate that the phosphorylation of the IDRs plays a critical role in the regulation of NPM1 oligomer stability.

**Figure 6.**
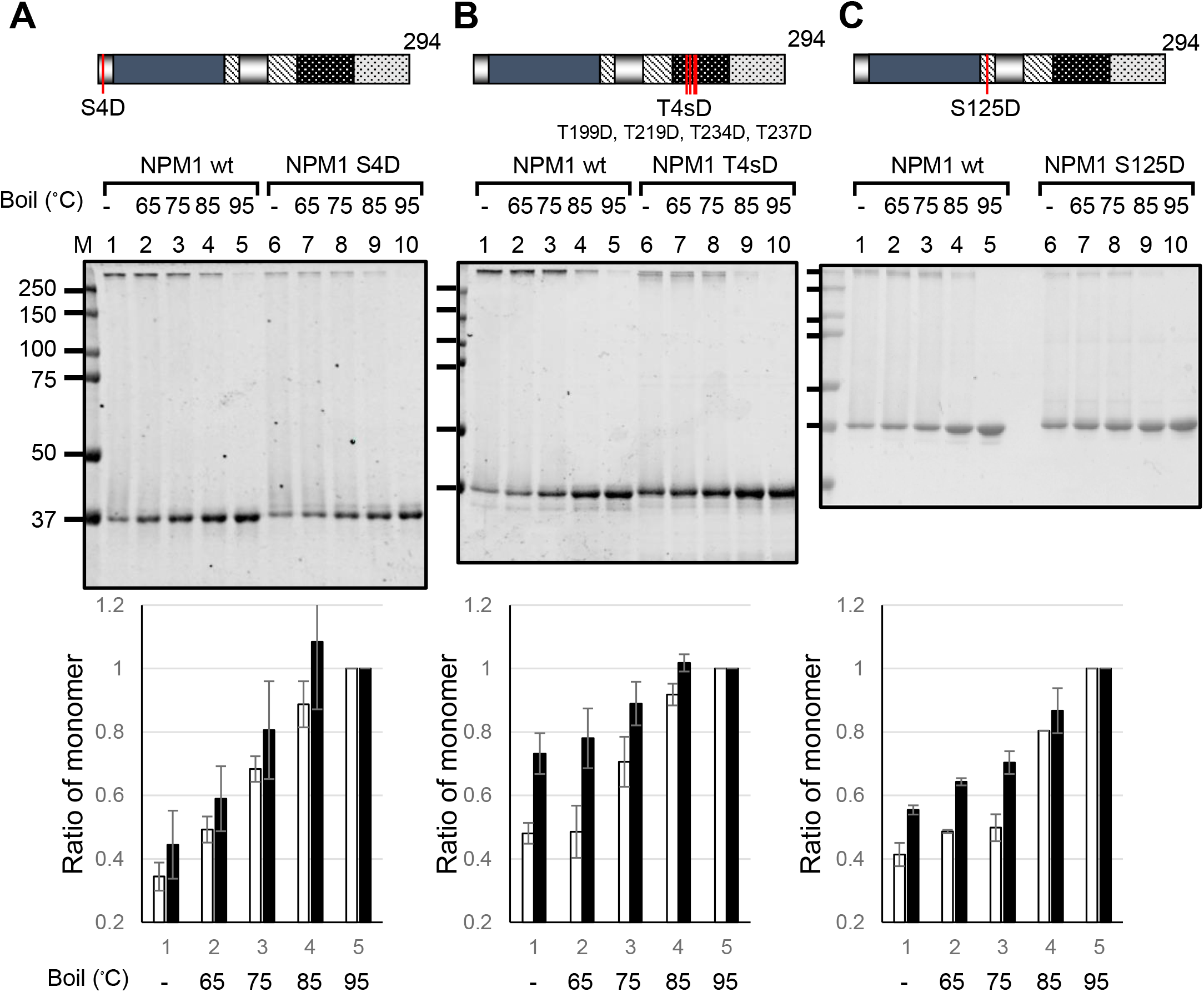
Effects of phosphomimetic mutations at the IDRs on the stability of NPM1 oligomers. NPM1-S4D (A), NPM1-T4sD (B), and NPM1-S125D (C) (500 ng) were separated by 10% SDS-PAGE before or after heating at 65°C, 75°C, 85°C, and 95°C for 5 min and visualized with CBB-staining. The band intensities of NPM1 monomers heated at 95°C in each experiment were set to 1.0 and relative intensities of monomers heated at different temperatures were measured by the Image J software and calculated (bottom graphs). White and black bars indicate wild-type and phosphomimetic mutants, respectively. The experiments were repeated three times and the data were averaged. Error bars indicate ±SD. Positions of mutations are shown at the top of the panels. Positions of molecular weight markers are shown at the left of the panels.

### Effects of NPM1 phosphorylation on its ability to induce LLPS

We next examined the effects of phosphorylation on the ability of NPM1 to form liquid droplets (Figures 7A–C). Liquid droplets were formed with both NPM1 wild-type and NPM1-S125D. In addition, both formed heterotypic liquid droplets efficiently with RNAs (Figures 7A and C). However, NPM1-T4sD formed small droplets or aggregates only at higher concentrations (Figure 7B) and droplets were not observed when RNAs were included. The inability of T4sD to form liquid droplets may be due to the inhibitory effects of bIDR phosphorylation on inter-oligomer interactions [12].

**Figure 7.**
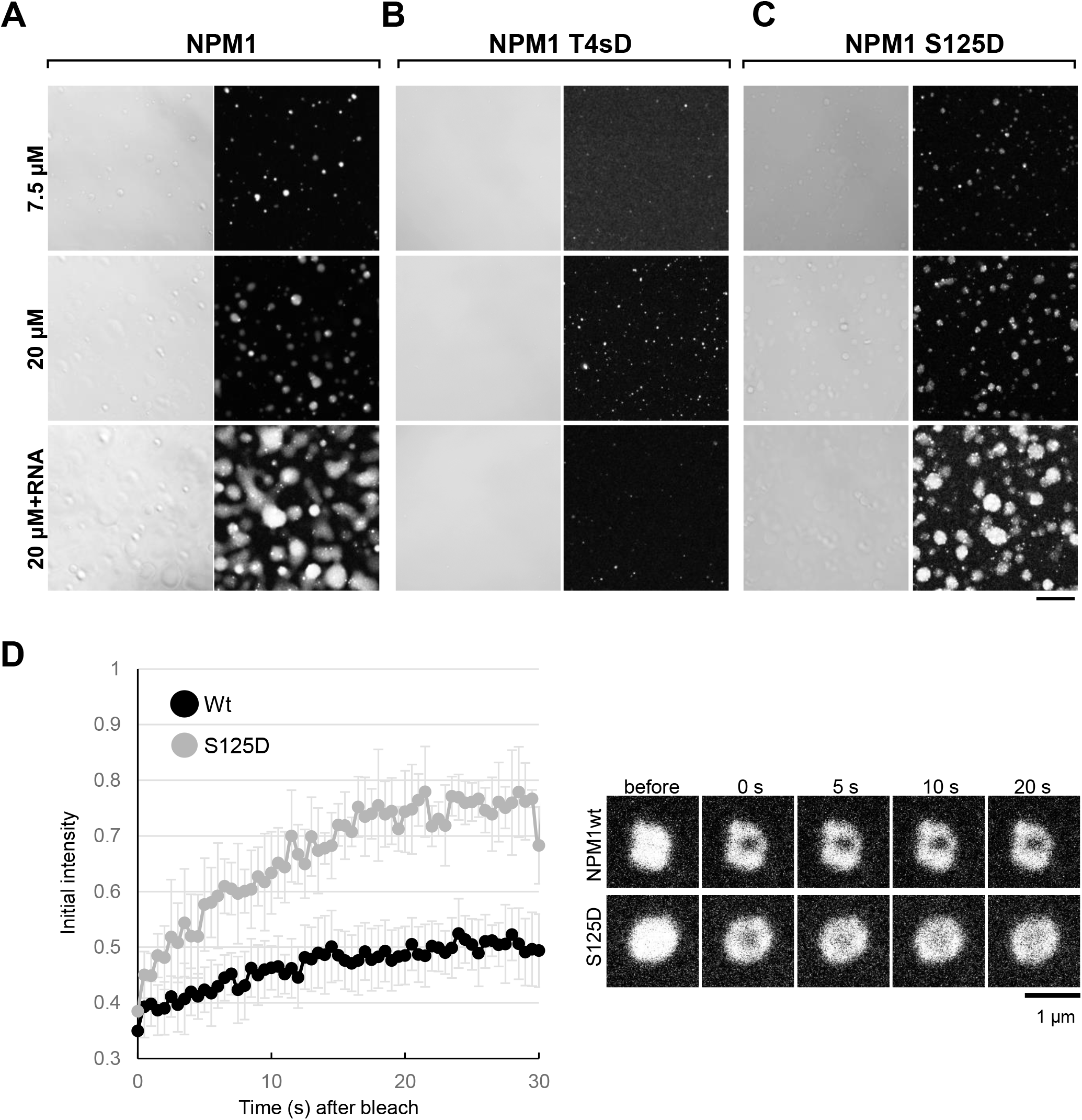
Liquid droplet formation by NPM1 phosphomimetic mutants. A–C. Liquid droplet formation by phosphomimetic mutants. Liquid droplets were formed with NPM1 (A), NPM1-T4sD (B), and NPM1-S125D (C) labeled with IC3-OSu in the buffer containing 2% Ficoll PM70 and 150 mM NaCl. The concentration of NPM1 was 7.5 or 20 μM and 1 μg RNAs were included in the bottom panels. Bright-field (left) and fluorescent images (right) are shown for each sample. A bar at the right bottom indicates 10 μm. D. FRAP analyses of NPM1 wild-type and S125D mutant. FRAP analyses were performed as in Figure 5C with NPM1 wild-type (black) and S125D (gray). For each sample, 10 droplets were photobleached and the data were averaged. Error bars indicate ±SD. Typical droplets before and 5, 10, and 20 sec after photobleaching with a 560 nm laser are shown at the right. A bar at the right bottom indicates 1 μm.

As shown in Figure 6, NPM1 oligomer stability is correlated with liquid droplet fluidity. The Ser125 in NPM1 is constitutively phosphorylated and the Ser125 phosphomimetic mutant exhibited lower oligomer stability (Figure 6C). We next examined the effect of phosphomimetic mutation at Ser125 on liquid droplet fluidity using a FRAP assay. The recovery rate of droplets formed by NPM1 wild-type was slow and the intensity of the bleached region recovered to only 50% of the initial intensity after 30 sec. Furthermore, the recovery rate for NPM1-S125D was far higher than that for the wild-type, and 70%–80% of the initial intensities were recovered after 30 sec. These results suggest that phosphorylation at IDRs regulates the stability of NPM1 oligomers and thereby determines the dynamic behavior of liquid droplets.

## Discussion

### Regulation of the oligomerization of NPM1 by acidic amino acids in the IDRs

In this study, we examined the effects of N-terminal and central acidic amino acids on the stability of NPM1 oligomers. MD simulations suggest that acidic IDRs move randomly with a minor preference for the oligomerization domain, which attenuates the monomer–monomer interactions. The interaction network formed with Tyr67 in one monomer and with Tyr17, His40, and His115 in the other monomer is likely to be disrupted by the acidc disordered regions. Any of the clustered acidic amino acids in the IDRs are capable of interacting dynamically with the oligomerization domain to disrupt the aforementioned interaction network. Thus, an increase in the net-negative charges in the acidic IDRs by phosphorylation decreases the stability of NPM1 oligomers. Consistent with this, phosphomimetic mutations at Ser4 and Ser125 in the acidic disordered regions decrease the stability of the oligomers, although the effect of phosphomimetic mutation at Ser4 on the stability of the oligomers was minimal. Phosphorylation at sites in the oligomerization domain was shown to decrease the stability of the oligomers [31]. Our data provide a novel mechanism in which the stability of NPM1 oligomers is regulated by acidc disordered regions and their phosphorylation.

### Regulation of oligomer stability and homotypic LLPS of NPM1

NPM1 undergoes LLPS [26, 27]. Previous studies have indicated that oligomerization and the inter-oligomer interactions that occur via the aIDR and the bIDR are required for the induction of LLPS. Although the effects of inter-oligomer interactions between the aIDR and the bIDR on the ability of NPM1 to promote LLPS have been analyzed [27], the effects of oligomer stability on the induction of LLPS have not been well characterized. Our results demonstrate that partial destabilization of NPM1 oligomers (such as that occurring with H40A, H115A, and S125D mutants) did not clearly affect the formation of liquid droplets; however, the fluidity of the droplets increases. This observation indicates that regulation of oligomer stability is an important mechanism in determining the fluidity of the nucleolus. At least three factors are thought to be involved in the regulation of oligomer stability and the ability to form liquid droplets with NPM1: 1) the translation initiation position, 2) post-translational modifications of the IDRs, and 3) post-translational modifications of the oligomerization domain itself.

We previously reported that interactions between the aIDR and the bIDR occur both inter- and intra-molecularly, and that the C-terminal globular domain that is required for RNA recognition attenuates the intra-molecular interactions, resulting in the formation of the inter-molecular interactions between oligomers [12]. Therefore, B23.2, a splicing variant of NPM1 that lacks a part of the C-terminal globular domain, was inefficient at forming liquid droplets (data not shown). Furthermore, the interactions between the aIDR and the bIDR are strongly inhibited by phosphorylation at Thr199, Thr219, Thr234, and Thr237, which introduces negative charges in the bIDR [12]. Inhibiting the inter-molecular interactions between the aIDR and the bIDR of NPM1 attenuates the process that drive LLPS. This is consistent with a recent report that decreasing ‘charge blockiness’ in the bIDR in NPM1 inhibits LLPS [32]. We also emphasize that the mitotic phosphorylation in the bIDR not only suppresses the inter-oligomer interactions, but also decreases the NPM1 monomer–monomer interactions. We propose that these two mechanisms synergistically lead to the disassembly of the nucleolus during mitosis.

It is currently unknown how much NPM1 is free from the other nucleolar components in living cells. NPM1 oligomers are suggested to be a scaffold of the GC region of the nucleolus that is formed by interactions with various Arg-rich motif proteins and pre-rRNAs bound to ribosomal proteins. NPM1-centered reversible homotypic and heterotypic interactions are likely to be important for the formation of a nucleolus with high fluidity. The high fluidity of the nucleolus may allow efficient formation and transport of ribosome precursors that are synthesized in the central region of the nucleolus. As previously reported, the homotypic liquid droplets that are formed by NPM1 alone are “aged” and the fluidity of the NPM1 droplets decreases with incubation time [27]. Therefore, we believe that a dynamic equilibrium between homotypic and heterotypic phase separation by NPM1, RNAs, and proteins containing Arg-rich regions is required to maintain the fluidity of the GC region and, thus, the integrity of the nucleolus.

## Methods

### Plasmid construction

The cDNAs for M5-NPM1 and M7-NPM1 were amplified by polymerase chain reactions (PCR) with primer sets, either 5’-agctagcatatggacatggacatgagccc-3’ or 5’-agctagcatatggacatgagccccctgag-3’ for M5 or M7, respectively, as forward primers and T7-terminator primer as a reverse primer, and pET14b-NPM1 as a template using KODplus Neo (Toyobo). For the construction of pET14b-NPM1-6A, -NPM1-23A, -NPM1- 236A, -NPM1-2346A, and -NPM1-234567A, forward primers, 5’-aaaaaacatatggaagattcgatggccatggacatgagccccctg-3’, 5’-aaaaaacatatggcagcttcgatggacatggacatga-3’, 5’-aaaaaacatatggcagcttcgatggccatggacatgagccccctg -3’, 5’-aaaaaaaacatatggcagcttcgatggccatggacatgagccccctg -3’, and 5’-aaaaaaaacatatggcagctgcggcggccgcggacatgagccccctgagg-3’, respectively, were used. For the construction of the C-terminal truncation mutants, NPM1(1–150), (1–140), (1– 130), and (1–117), a forward primer (T7 promoter primer) and reverse primers, 5’-aaaaggatccttacttgctaccacctccagggg 3’, 5’-aaaaggatccttatccagatatacttaagagtt-3’, 5’-aaaaggatccttactcctcttcatcttctgact-3’, and 5’-aaaaggatccttatactaagtgctgtccactaat-3’ respectively, were used. For NPM1-K32A, -Y67H, -Y67F, -H40A, -H115A, two step PCR was performed to amplify the mutant cDNAs using primer sets, 5’-cactttgcggtggataatgatgaaaatg-3’ and 5’-atccaccgcaaagtgataatctttgtcgg-3’, 5’-gcaatgaatcacgaaggcagtccaattaaa-3’ and 5’-actgccttcgtgattcattgcctctgcttc-3’, 5’-caatgaatttcgaaggcagtccaattaaag-3’ and 5’-actgccttcgaaattcattgcctctgcttc-3’, 5’-aaaatgaggcccagttatctttaagaacggt-3’ and 5’-aagataactgggcctcattttcatcatta-3’, and 5’-gtggacaggccttagtagctgtggaggaag-3’ and 5’-gctactaaggcctgtccactaatatgcactg-3’, respectively. The amplified cDNAs were digested with Nde I and BamH I, and cloned into the same sites of pET14b. Plasmids for wild-type and other mutant NPM1 were previously described [2, 33].

### Protein expression and purification

The *E. coli* BL21 (DE3) strain that was transformed with pET14b vectors was grown in 300 ml LB media containing ampicillin (0.1 mg/ml) at 37°C until OD_600_ reached 0.5– 0.7, and protein expression was induced by the addition of isopropyl β-D-thiogalactopyranoside (0.9 mM). The cells were incubated for an additional 2 h, harvested by centrifuge at 7,000 rpm for 10 min, suspended in 10 ml of NPI-10 (50 mM NaH_2_PO_4_, 300 mM NaCl, and 10 mM imidazole [pH 8.0]) supplemented with 1 mM PMSF, sonicated for 30 s, three times, and centrifuged at 12,000 rpm for 15 min. The supernatants were mixed with Ni-NTA Superflow resin (100 μl) (QIAGEN) and incubated at 4°C overnight. The mixtures were loaded on a plastic column and the resin was washed with 10 ml of NPI-20 (50 mM NaH_2_PO_4_, 300 mM NaCl, and 20 mM imidazole [pH 8.0]) containing 1 mM PMSF. The bound proteins were eluted with NPI-250 (50 mM NaH_2_PO_4_, 300 mM NaCl, and 250 mM imidazole [pH 8.0]). The peak fractions were dialyzed with buffer H (20 mM Hepes pH7.9, 10% glycerol, 1 mM PMSF) containing 500 mM NaCl at 4°C for more than 3 h. The dialyzed proteins were kept at −80°C until use. To label the purified proteins, His-tagged proteins bound to Ni-NTA resins were incubated with IC3-OSu dye (DOJINDO) in NPI-20 buffer at 4°C for 1 h. The resin bound by His-NPM1 proteins was washed and the proteins were eluted as above. For oligomer detection assays shown in Figure 1, the purified NPM1 proteins were separated by 10 % SDS-PAGE, cut out, and eluted from the gel, followed by the denature-renature protocol as described previously [12].

### Liquid droplet formation and FRAP assay

Liquid droplet formation assay was performed according to the methods described previously [34]. Briefly, the salt (NaCl) concentration of the proteins in buffer H containing 500 mM NaCl was diluted to 150 mM and the samples were supplemented with 4% Ficoll PM70 (Sigma-Aldrich). Then the samples were incubated in a glass-based 384-well plate (EZVIEW assay plate, IWAKI glass) at room temperature. Droplets were observed within 1 h after sample preparation under a confocal microscope with a 63×/1.4 oil immersion-objective lens (LSM710, Carl Zeiss).

FRAP assay was performed with a confocal microscope as above. The central regions of fluorescent liquid droplets were bleached with 100% transmission of a 561 nm-laser and the images (512×512 pixels, zoom 15, scan speed 13, and a 561 nm-laser 0.5 %) were collected. The fluorescent intensities of the bleached area (Ia), a non-bleached area in the same droplets (Ib), and the area outside of the droplets (Ic) were measured every 0.5 s for 30 s. The relative fluorescence intensities (RFI) were calculated as following: RFI = (Ia-Ic) / (Ia0-Ib0) × (Ib0-Ic0) / (Ib-Ic). The initial RFI was set to 1.0 and the relative intensities of at least 10 experiments were averaged.

### Computational analysis

The crystal structure of NPM1 pentamer (amino acids 14–118, PDB code: 5EHD) was preprocessed for the computational analyses using Protein Preparation Wizard [35] in the Maestro program of Schrödinger Suite 2019-1 (Schrödinger, LLC, New York, USA), which added hydrogen atoms, repaired the side chains of incomplete residues, and optimized the hydrogen-bonding network. All crystallographic water molecules were removed, and then the restrained minimization of protein structure was performed with an RMSD cut-off value of 0.3 Å using the OPLS3e force field. The missing N-terminal and central acidic regions were compensated using the Prime program [36, 37] in Schrödinger Suite 2019-1, generating two structures NPM1(1–118) and NPM1(14–130), respectively. Three initial structures of NPM1 pentamer, NPM1(1–118), NPM1(14–130), and NPM1(14–118) were then subjected to the MD simulations. The simulations presented here used OPLS3e force field with explicit solvent and were run using the default parameters in Desmond program [38] of Schrödinger Suite 2019-1. The SPC water model was used. The systems were neutralized with Na^+^ or Cl^-^ ions and 0.15 M NaCl was also added. Periodic boundary conditions and a 9.0 Å cut-off for non-bonding interactions were used with electrostatic interactions treated using the particle mesh Ewald method with a tolerance of 10^−9^. After a Desmond default relaxation protocol, 100 ns simulations in a normal pressure and temperature ensemble (temperature: 300 K, thermostat relaxation time: 1.0 ps, pressure: 1 atm, and barostat relaxation time: 2.0 ps) were performed using a Nose-Hoover thermostat and a Martyna-Tobias-Klein barostat. Trajectory atomic coordinate data were recorded every 20 ps. Five simulations for each initial structure of NPM1 pentamer were repeated by varying the random seeds for generating the initial velocities of the atoms. From the last 50 ns trajectory of each simulation, 2,500 structures of NPM1 pentamer (in total 12,500 structures for each initial structure) were obtained. Using these structures, the number of the contacts (hydrogen-bonding, ionic, π-π stacking, and π-cation interactions) between the oligomerization domain and the aIDRs (N-terminal region in NPM1 (1–118) and the central acidic region in NPM1 (14–130)) were enumerated by Simulation Interactions Diagram analyses in Desmond program (Schrödinger Suite 2019-1). The number of monomer–monomer contacts in the oligomerization domain was also enumerated in NPM1 (1–118), NPM1 (14–130), and NPM1 (14–118).

## Author contributions

MO, SE, NJ, TU, KN, and TK performed biochemical experiments and analyzed data. SIO and NT performed MD simulations and analyzed data. MO, SIO, and TN designed and supervised the research. MO and SIO wrote the manuscript, and all authors contributed to editing the manuscript.

### Conflict of interest statement

None declared.

## Acknowledgment

We thank Ms Hirawake-Mogi for plasmid preparation. We also thank Enago for the English language review. This work was supported by KAKENHI Grants from the Ministry of Education, Culture, Sports, Science and Technology of Japan to MO (17K07300).

## Abbreviations

LLPS: liquid-liquid phase separation
NPM1: nucleophosmin 1
IDR: intrinsically disordered region
MD simulation: molecular dynamics simulation
rRNA: ribosomal RNA
FRAP: fluorescence recovery after photobleaching
GC: granular component

## Notes

### Competing Interest Statement

The authors have declared no competing interest.

